# Confidence modulates exploration and exploitation in value-based learning

**DOI:** 10.1101/236026

**Authors:** Annika Boldt, Charles Blundell, Benedetto De Martino

## Abstract

Uncertainty is ubiquitous in cognitive processing, which is why agents require a precise handle on how to deal with the noise inherent in their mental operations. Previous research suggests that people possess a remarkable ability to track and report uncertainty, often in the form of confidence judgments. Here, we argue that humans use uncertainty inherent in their representations of value beliefs to arbitrate between exploration and exploitation. Such uncertainty is reflected in explicit confidence judgments. Using a novel variant of a multi-armed bandit paradigm, we studied how beliefs were formed and how uncertainty in the encoding of these value beliefs (belief confidence) evolved over time. We found that people used uncertainty to arbitrate between exploration and exploitation, reflected in a higher tendency towards exploration when their confidence in their value representations was low. We furthermore found that value uncertainty can be linked to frameworks of metacognition in decision making in two ways. First, belief confidence drives decision confidence—that is people’s evaluation of their own choices. Second, individuals with higher metacognitive insight into their choices were also better at tracing the uncertainty in their environment. Together, these findings argue that such uncertainty representations play a key role in the context of cognitive control.

## Introduction

All cognitive computations are plagued by uncertainty (Bach & Dolan, 2012). On the one hand, uncertainty can be inflicted on our cognitive systems through external sources, for example when we perceive noisy information in the environment. On the other hand, uncertainty can arise directly from the way the brain processes information. Recent research has supported the view of a signal-inherent representation of uncertainty, suggesting a coding scheme in which the reliability of a signal is represented together with its average strength (e.g., Ma, Beck, Latham, & Pouget, 2006; Ma & Jazayeri, 2014; Van Bergen, Ma, & Pratte, 2015). Such a coding scheme allows the decision maker to more efficiently integrate new evidence by giving more weight to a reliable evidence source and discounting information from a source identified as unreliable. While such flexible weighting of evidence might take place automatically and without agents being aware of it, recent findings suggest that humans can accurately trace uncertainty and report the level of confidence in their judgments (Shea, Boldt, Bang, Yeung, Heyes, & Frith, 2014; Meyniel, Sigman, & Mainen, 2015b; Guggenmos, Wilbertz, Hebart, & Sterzer, 2016).

Most confidence studies focus on only the last stage of the decision-making process, with confidence reflecting the internal belief as to whether the chosen option was the ‘good’ or ‘correct’ one (e.g., Pleskac & Busemeyer, 2010; Baranski & Petrusic, 1994; Fleming, Weil, Nagy, Dolan, & Rees, 2010). More recently, Pouget and colleagues (Pouget, Drugowitsch, Kepecs, 2016) proposed that we should distinguish decision confidence from certainty. While the former tracks the probability of an action to be correct, the latter indexes the uncertainty in a representation. Following a similar approach with this study, we separately measure the evolution of these two quantities during learning. We define decision confidence as the confidence that an action (e.g. choosing the most valuable bandit) is correct. We define belief confidence as the agent’s internal readout of the uncertainty inherent to her belief about the value of each bandit, related to the concept of certainty proposed in the framework by Pouget and colleagues (Pouget, Drugowitsch, & Kepecs, 2016). More specifically, when people assign a value to an object on which they base their preferences and choices, those beliefs can vary with regard to how precise they are: on our first day at work, we might guess that we will like our new job, but several months later our certainty in that belief might have increased, resulting in a highly precise value belief representation. While recent studies have started to investigate decision confidence (in the context of value-based choice; e.g. De Martino et al., 2013) and belief confidence (Lebreton et al., 2015; De Martino, Bobadilla-Suarez, Nouguchi, Sharot, & Love, 2017), the interplay between these two types of confidence has been understudied.

Using a novel variant of a multi-armed bandit task, we show that we can reliably measure how confidence in people’s beliefs evolves over the course of learning. The aim of our study was twofold: to study the link between these two types of confidence and to understand how people use their belief confidence to guide decision making. First, we investigate the links between people’s belief confidence and decision confidence. We have previously shown that confidence in value-based choices is largely driven by the difference in values between choice options (De Martino, Fleming, Garrett, & Dolan, 2013) and that such confidence is related to changes of mind in future value-based decisions (Folke, Jacobsen, Fleming, & De Martino, 2016). Here, we furthermore show that belief confidence contributes to decision confidence. In other words, the level of certainty in the value we assigned to something can increase our decision confidence.

The second goal of our study was to investigate how an agent uses the explicit representations of the uncertainty inherent in value representations (belief confidence) to arbitrate between different behavioral strategies. Previous research has highlighted the role of confidence signals as internal teaching signals that support cognitive control (Fernandez-Duque, Baird, & Posner, 2000), learning (Guggenmos et al., 2016), and social interactions (Bahrami, Olsen, Latham, Roepstorff, Rees, & Frith, 2010). Here, we are focusing on higher-order action selection, the so-called exploration-exploitation tradeoff (Schumpeter, 1934; Sutton & Barto, 1998; Cohen, McClure, & Yu, 2007; Kolling, Behrens, Mars, & Rushworth, 2012). Consider, for instance, that when choosing a restaurant for dinner, you must decide whether to go to your usual local restaurant (exploitation) or try a new restaurant that has just opened down the road (exploration)—hoping to find a new favorite while accepting the risk of consuming a bad meal. Arbitrating optimally between exploration and exploitation is not trivial and different algorithms for achieving a good balance between these extreme behavioral strategies have been discussed in the machine learning literature. A principled and efficient way of arbitrating between these modes is to deploy exploration towards the options or actions regarding which the agent is more uncertain (Dayan and Sejnowski, 1996; Gittins and Jones, 1974). Experimental work has shown that people can implement such sophisticated strategies, which take into account the level of uncertainty in their environment (Daw, O’Doherty, Dayan, Seymour, & Dolan, 2006; Frank, Doll, Oas-Terpstra, & Moreno, 2009), even if there are substantial inter-individual differences (Badre, Doll, Long, & Frank, 2012).

These studies have deployed computational modelling as a tool to estimate the amount of uncertainty inherent in the belief representation, which then in turn has been shown to affect subsequent choices. However, direct measurement of uncertainty in the form of confidence judgments has to date not been attempted. Here we show how confidence guides the trade-off between exploration and exploitation. We moreover aimed to answer the question of whether the degree by which people can accurately harness the level of uncertainty in their beliefs through confidence relates to their metacognitive accuracy (the degree to which their perceived accuracy corresponds to their objective accuracy) might explain some of the inter-individual variability reported in the literature. Here we present the results from two studies in which participants continuously faced two lotteries (two-armed bandits), each associated with a different average outcome. The participants’ task was to maximize their earnings by choosing the most rewarding bandit. Furthermore, participants had to rate the value of the lotteries together with the confidence they held in this value belief (belief confidence), which we predicted would guide their choices. From time to time, participants were furthermore asked to freely choose between the two lotteries and to rate how accurate they regarded their choice (decision confidence). The first experiment focused on the development of preferences and confidence judgments over time. The second study focused on the decision stage and the influence of belief confidence on the trade-off between exploration and exploitation. Taken together, our findings argue that the explicit representation of uncertainty expressed through confidence report plays a key role in arbitrating between complex decision strategies.

## 1. Methods and Materials

### 1.1 Participants

In Experiment 1, we tested 22 participants, 20 of which were female, and 3 were left-handed. Participants’ ages ranged from 18 to 32 years (M = 24.0). One participant had to be excluded because during debriefing she reported that she had tracked the value of the bandits with a piece of paper. The final sample therefore included 21 participants. For Experiment 2, we tested 30 participants, 15 of whom were female, and 1 was left-handed. The participants’ ages from Experiment 2 ranged from 19 to 34 (M = 26.4). No participants were excluded in this experiment.

For both experiments, all participants reported normal or corrected-to-normal vision, were English-proficient, and reported that they had no psychiatric or neurological disorders or gambling addiction. All participants gave informed consent and were reimbursed for their time (*£*10/hour). Each session lasted approximately 90 minutes, including instruction and debriefing. In addition, participants also received performance-dependent rewards (Experiment 1: M = *£*8.82; min = *£*8.44; max = *£*8.97; Experiment 2: M = *£*8.71; min = *£*8.17; max = *£*9.11). All procedures were approved by the local ethics committee.

### 1.2 Task and Procedures

Experiment 1 comprised 75% rating trials and 25% choice trials. During rating trials, participants were first presented with the outcome of one of the randomly chosen arms of the bandit. They then had to rate the value belief associated with this arm (average number of points that could be won from this machine) and their confidence in this estimate. We used a two-dimensional grid scale to collect these measurements. Value-belief judgments ranged from 0 to 100 points, belief confidence judgments from “guessing” to “very confident”. Participants were told that during rating trials, they would observe another person gamble at one of the two slot machines. Unbeknownst to the participant, this person chose one of the slot machines randomly with equal probability. During choice trials, participants could freely choose one of the two arms of the bandit and were then asked to rate their confidence in being correct in their choice before seeing the outcome of the trial. Decision confidence ratings were given on a scale that ranged from “guessing” to “very confident”. Points won on those trials contributed towards the final reimbursement sum participants received after the experiment and they were instructed to carefully use the information previously gained when making their choices. Both trial types were intermixed randomly. Figure 1 shows an example of three trials (2 rating and 1 choice trial).

**Figure 1.**
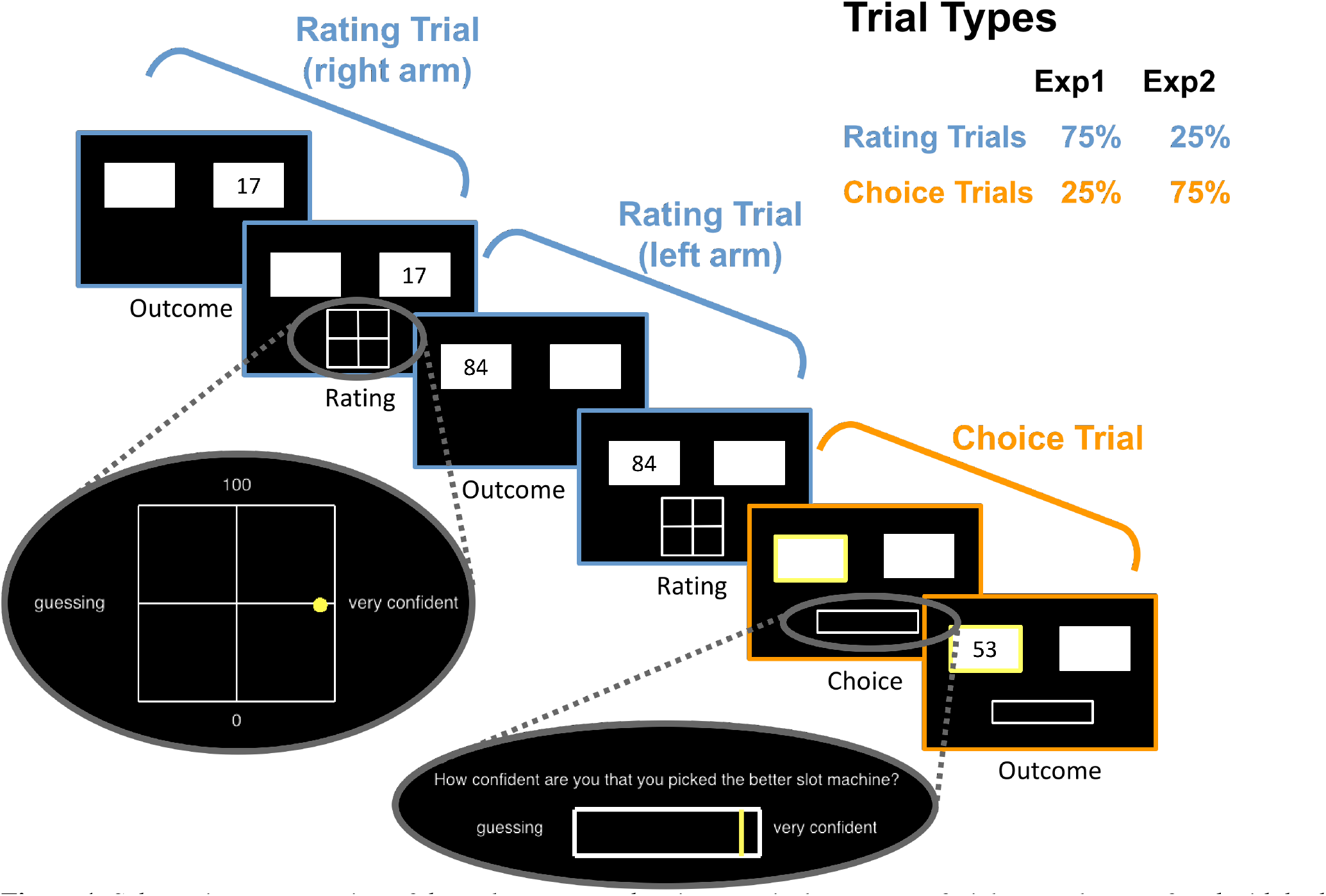
Schematic representation of the task structure, showing a typical sequence of trials: People were faced with both rating (blue) and choice (orange) trials. During rating trials, they observed outcomes randomly from one arm of the 2-armed bandit, represented as squares. Participants then rated the average value of this arm and their confidence in this value-belief estimate on a two-dimensional grid. During choice trials, participants freely chose one arm of the bandit, rated their confidence in this decision and were then shown the reward. In Experiment 1, 75% of trials were rating trials and 25% trials were choice trials, with both trial types intermixed randomly. In Experiment 2, these proportions were reversed.

In total, participants completed 600 trials, grouped into blocks of different lengths (20 to 60 trials long). Participants were thus unable to predict how many evidence samples they would be able to collect in the present block, allowing us to avoid the possibility that they would adjust their risk-taking behavior given how close they were to the end of the block (Kolling, Wittmann, & Rushworth, 2014). For each block, two different beta distributions [(a = 1; b = 3), (a = 2; b = 3), (a = 3; b = 3), (a = 3; b = 2), (a = 3; b = 1)] were taken to generate the rewards of the two arms of the bandit. These parameter ranges were chosen so that the resulting distributions had different skews away from uniform whilst being unimodal. The resulting samples were multiplied by 100 to achieve rewards bounded between 0 and 100 points. Prior to the experiment, participants completed three practice blocks, introducing them to choice and rating trials separately (5 trials each) and together (20 trials). Importantly, all participants completed exactly the same blocks, shuffled with regard to the order in which they appeared. The gamble outcomes that participants were presented with were thus the same for everyone, only dependent on their choices.

A key question we aimed to address with Experiment 2 was whether people would use their belief confidence to arbitrate between exploration and exploitation. While the high number of rating trials (75%) in Experiment 1 allowed us to assess whether such belief confidence was related to the learning process in a meaningful manner, this design was ill-suited to investigate how people chose between exploration and exploitation. This is primarily due to the high proportion of rating trials, in which participants observed outcomes from both arms of the bandit, which allowed them to form a good enough estimate of the value of each arm of the bandit and making exploration unnecessary. As a result, their choices should arguably mostly be exploitative. In Experiment 2, which was highly similar to Experiment 1, we therefore reversed the ratio of trial types with most trials (75%) being choice trials. However, to allow insight into the evolution of value estimation during learning, we also collected a smaller percentage (25%) of rating trials, but for the reason mentioned above in this new version we no longer let participants observe the outcomes of these trials. During rating trials, people now judged their value beliefs and belief confidence for both lotteries. Which bandit had to be rated first was randomly chosen by the computer and could not be predicted by the participants. With a considerably larger number of choice trails, we expected more exploration trials in this experiment, that is trials in which people would not just classify their confidence as “guessing” if asked, but instead knew that they had chosen the lower-value option. The confidence scale was thus extended to range from “low confidence” to “high confidence” for this experiment. Prior to Experiment 2, participants completed two practice blocks, familiarizing them first with choice trials (5 trials) and then with a combination of both choice and rating trials (12 trials).

All testing was administered using a 24-inch monitor (16:9 aspect ratio) using the MATLAB toolbox PTB3. Responses were given with an ordinary computer mouse. Prior to the analyses reported in this study, we excluded on average 2.2% of choice trials from the analyses of Experiment 1. Those were trials in which participants responded too slowly (+/− 3 σ rule; min = 0.7%; max = 3.3%). The same held for 1.7% of choice trials in Experiment 2 (min = 0.2%; max = 2.9%). No rating trials were excluded.

### 1.3 Data Analyses

The first set of analyses for Experiment 1 aimed to assess whether our new paradigm accurately captured belief learning over time and whether the uncertainty inherent in those beliefs — our newly introduced concept belief confidence — constituted a meaningful judgment. We therefore attempted to link belief confidence to the more traditionally used concept of decision confidence using a linear, hierarchical regression model to predict decision confidence. We included participants’ last rated value-belief and belief-confidence ratings of both the chosen and the unchosen option as fixed effects, as well as the interaction of these estimates. Moreover, three control variables were included as fixed effects. The first variable was the objective accuracy of a trial. Decision confidence is an agent’s subjectively perceived accuracy of being correct. The objective accuracy of a trial should thus positively predict confidence, with higher confidence for correct trials. The second control variable were log-transformed RTs, given the time heuristic that suggests that RTs and confidence are negatively correlated across trials (e.g. Audley, 1960; Moreno-Bote, 2010; Hanks, Mazurek, Kiani, Hopp, & Shadlen, 2011; Kiani, Corthell, & Shadlen, 2014). The third and last control variable was the number of the current trial within the block. We predicted that people’s decision confidence would increase over the course of each block, reflecting increasingly better choices. In addition to these fixed effects, the data were modeled with a random intercept and random slopes for all regressors. This regression model was fit to data from choice trials only with the R package *lme4*. We used Satterthwaite approximations (Schaalje, Brigham, & Fellingham, 2001) to obtain degrees of freedom and to calculate p-value estimates. All predictors were z-transformed prior to being entered into the model to obtain standardized regression coefficients.

The key goal of Experiment 2 was to test for uncertainty-driven exploration. Here, we define exploration as choice trials in which participants chose the lower value option. Since value ratings for the individual arms of the bandit were only collected during rating trials, we extrapolated the current value rating for choice trials using the measurement taken at the most recent rating trial. An exploration trial is then defined as a trial in which the participant choses the arm of the bandit they rated as yielding lower rewards, whereas an exploitation trial is defined as choosing the option rated to yield higher values. We then fitted a logistic, hierarchical regression model to the choice-trial data of Experiment 2 to test for uncertainty-driven exploration. The dependent, binary variable expressed whether or not on the current trial people chose the choice option that they had previously rated as having a lower value compared to the other option—our new operationalization of exploration. We included participants’ last rated belief confidence of both the higher- and the lower-value choice option as fixed effects, as well as the interaction of these estimates. Moreover, the difference in value for the choice options (high minus low) was included as a fixed effect. In addition to these fixed effects, the data were modelled with a random intercept and random slopes for all regressors. This regression model was again fit to data from choice trials only, using the R package *lme4*. All predictors were z-transformed prior to being entered into the model to obtain standardized regression coefficients.

In Experiment 2, we furthermore attempted to link interindividual differences in people’s ability to trace uncertainty to their metacognitive efficiency. We obtained metacognitive efficiency measures, M-ratio, by fitting second-order signal-detection theory (SDT) models to participants’ decision confidence data (Maniscalco & Lau, 2012; Fleming & Lau, 2014) using the MATLAB package HMeta-d (Fleming, 2017). This approach automatically adjusts for differences in first-order performance. We furthermore fitted hierarchical, linear regression models that predicted belief confidence from the variance and mean of past outcomes, the current trial within the block, as well as the arm of the bandit. The standardized beta weights of the influence of the variance of past outcomes on belief confidence were used as an indicator for the extent to which people were capable of tracking uncertainty.

For the sake of simplicity, for all regression approaches in this study we report coefficients from the model that fitted the data best. However, a comprehensive list of models of varying degrees of complexity (10, 5, or 6 models respectively) and a formal model comparison based on BIC scores is included the Supplementary Material.

## 2. Results

### 2.1 Experiment 1

#### 2.1.1 Participants formed value beliefs over time

In Experiment 1, we investigated how value belief and belief confidence rating evolved over time. We assumed that participants would adjust their value-belief estimates with each sample of evidence, presumably getting closer to the true value over time. At the same time, the level of confidence in these value estimates should increase in the course of learning. We found support for both of these predictions and address each in turn here. First, as expected we found that participants’ value estimates increased in accuracy over time. Value estimation accuracy was on average 11 points off during the first half of the blocks, and only 8 points off during the second half. This difference in estimation errors was reliable, t(20) = 9.7, p <0.001. This overall pattern of improvement can furthermore be seen in Figure 2A, in which the grey hairline arrows are smaller for later trials in the example block (darker colors) compared to earlier trials (lighter colors). This figure furthermore shows that people perceived the arms of the bandit as more similar at the beginning of the block (the two traces are closer together) compared to the end of the block. Second, participants grew monotonically more confident in their value estimates (belief confidence) over the course of learning, M_earty_ = 0.36 vs. M_late_ = 0.58, t(20) = 9.4, p <0.001 (see Figure 2B). This increase in confidence is furthermore reflected in the traces in Figure 2A, where later judgments (darker colors) lie closer towards the right end of the x-axis. However, the example traces also show that belief confidence could sometimes suddenly decrease over the course of a block, as reflected, for instance, in the two data points highlighted by asterisks in Figure 2A: the darker colored asterisk lies further to the left compared to the lighter colored asterisk. Presumably, this happens whenever the newly sampled evidence leads to a larger update in value belief. Indeed, the larger the absolute difference between the currently observed outcome and the previously estimated value for the respective arm of the bandit, the larger the decrease in belief confidence, as reflected in individual correlations that were negative for 20 out of 21 participants, rs >= −0.46 & rs <= −0.01, and reliable for 16 out of 21 participants (ps <0.01). Taken together, these findings suggest the paradigm used here is well suited to study the development of value beliefs and belief confidence over time: Participants’ value estimates increased in accuracy and this was reflected in increases in belief confidence.

**Figure 2.**
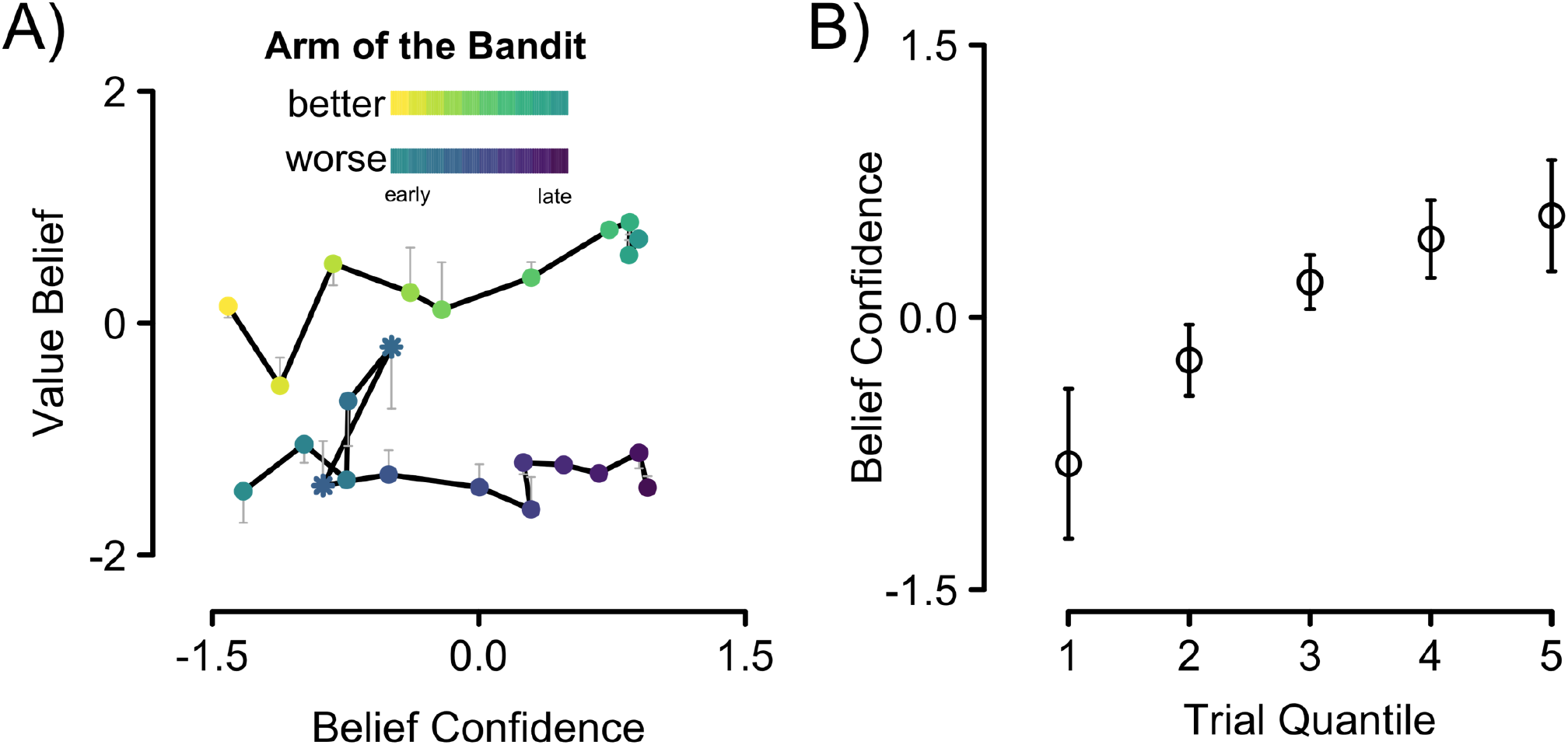
A) Average traces of participants’ value belief and belief confidence ratings, given on the 2-dimensional grid scale for one example block. All ratings are z-transformed within-subject and then averaged across participants to reduce inter-individual differences in the use of the rating scales. The arm with the objectively higher rewards is shown in yellow to green (shown on 10 trials out of the block, corresponding to one data point each) and the other arm in blue to purple hues (shown on 14 trials out of the block, corresponding to one data point each). The brightness reflects the position of the data points within with block with brighter (yellow or blue) hues representing the earlier trials. The hairline arrows reflect the mean reward, calculated from the observed outcomes (objective mean value of the past outcomes). The length of the arrows is therefore proportional to the estimation error with longer arrows reflecting worse value estimates. B) Belief confidence increased over blocks: The x-axis shows trial quintiles, calculated within each block ranging from the first (1) to the last (5) fifth of trials in each block. Belief confidence was z-transformed within-subject and then averaged across participants. The error bars reflect +/− 1 standard error of the mean.

#### 2.1.2 Linking belief confidence to decision confidence

Having established that our paradigm reflects meaningful changes in beliefs over time and uncertainty inherent in those beliefs, we then aimed to investigate the link between people’s belief confidence—the new concept introduced here—and decision confidence. We have previously shown that decision confidence is a function of the difference in value (De Martino et al., 2013). The schematic diagram in Figure 3A outlines this finding: two noisy value beliefs are represented as distributions, with their distance reflecting the difference in value (DV). In the present study, we furthermore propose that belief confidence affects decision confidence. Belief confidence is reflected in the spreads of the two belief distributions; the narrower these distributions are, the more confident people judged their decisions. We therefore fitted a hierarchical, linear regression model to predict people’s decision confidence from their value-belief and belief-confidence ratings of both the chosen and unchosen arm of the bandit, the interaction of these estimates, and three control variables: the objective accuracy of a trial and log-transformed RTs. The standardized regression coefficients for the fixed effects are presented in Figure 3B. We chose to present the best-fitting model here based on *BIC* scores (*BIC* = 5293.9). However, a detailed model comparison approach which included a range of models, both more parsimonious and more complex, is included in the Supplementary Materials. Some of these alternative models included the position of the trial within a block as a regressor (log-transformed).

**Figure 3.**
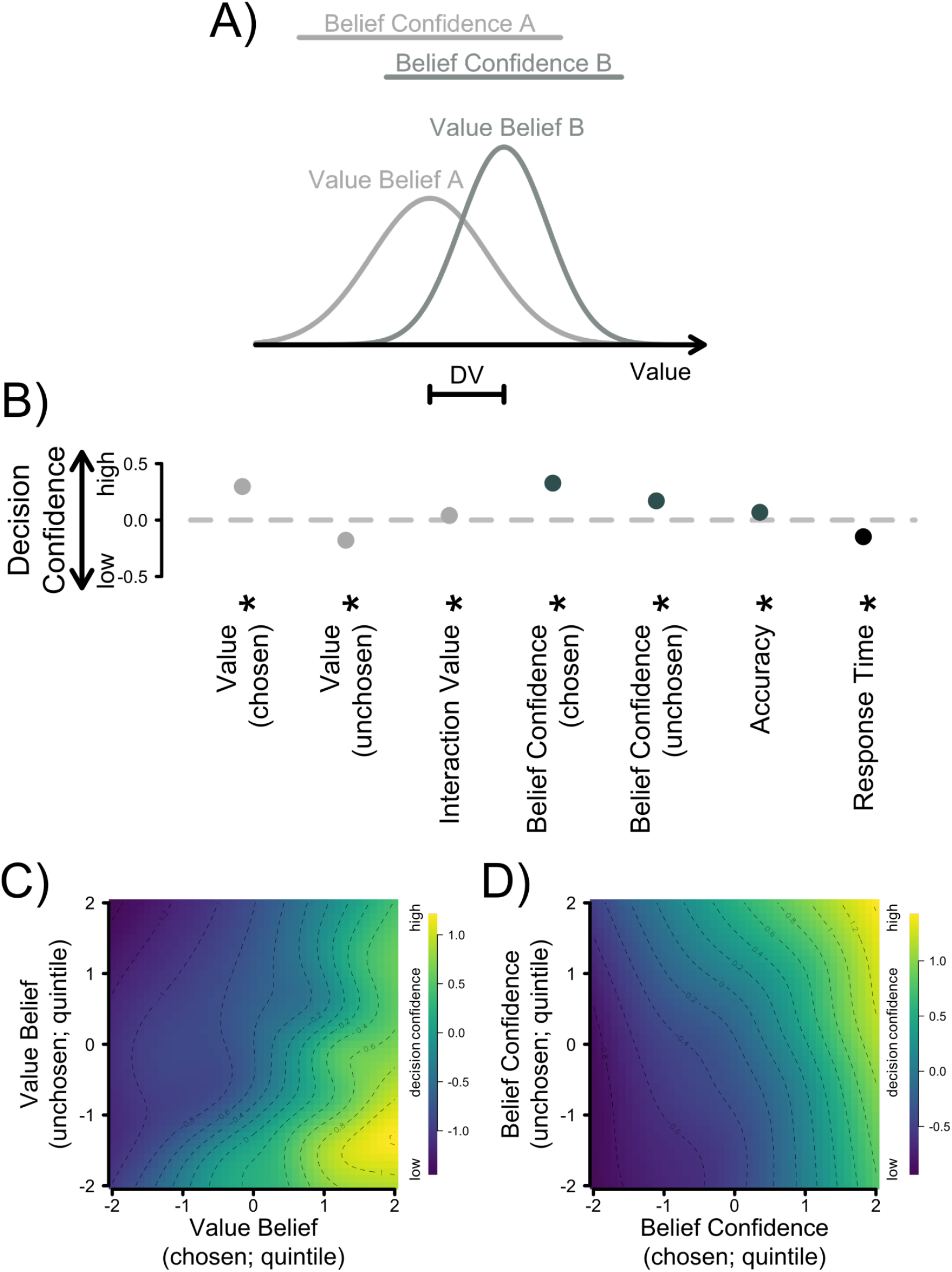
Hierarchical regression model used to predict decision confidence. A) Schematic figure showing the noisy value representation for two objects. For the purpose of simplicity, each value belief is represented as a normal distribution with a mean (value belief) and a standard deviation (belief confidence). For these two overlapping choice options, Option B has a higher value than Option A, and also a more precise value representation (higher belief confidence). DV = difference in value. B) Standardized, fixed regression coefficients from a hierarchical, linear regression model. Positive, higher parameter estimates reflect that an increment in this variable led to an increment in decision confidence. The error bars, which are almost entirely hidden behind the disks, reflect +/− 1 standard error of the mean. The light grey disks represent predictors linked to value, the dark grey disks represent predictors linked to belief confidence, and the black disks represent control variables. C) and D) depict the influence of the key predictors on decision confidence for both value belief (C) and belief confidence (D). Lighter colors reflect higher levels of decision confidence.

As reported by De Martino et al. (2013), a larger difference in value was associated with higher decision confidence, as reflected in the significantly positive and negative regression weights for the value of the chosen, *β* = 0.30, p <0.001, and unchosen option, *β* = −0.18, p <0.001, respectively. These predictors furthermore showed a small but reliable interaction effect, *β* = 0.04, p <0.05. Figure 3C depicts the influence of both value-belief regressors on decision confidence as a two-dimensional grid with lighter colors reflecting higher simulated decision confidence.

Critically, being confident in their value estimates also made people more confident in their decisions, both for the chosen, *β* = 0.33, p <0.001, and also the unchosen choice option, *β* = 0.17, p <0.001. The best-fitting model did not include an interaction between these two predictors. However, the prediction pattern formed by these two belief-confidence regressors is shown in Figure 3D.

Moreover, two control variables were included in these regression models. First, the objective accuracy of the current trial was a binary variable that affected decision confidence positively, *β* = 0.07, p <0.001, as predicted. In other words, if participants picked the objectively higher-value option, they tended to be more confident in their choices. Second, the faster a decision, the higher people’s subjectively-judged decision confidence, *β* = −0.15, p <0.001.

Taken together, the results of Experiment 1 suggest that our experimental setup allowed us to track the evolution of value beliefs over the course of learning and that belief confidence reflects meaningful insight into such learning. We moreover found that how certain people are in their value beliefs affected their decision confidence, in support of our first hypothesis.

### 2.2 Experiment 2

Having established with the findings from Experiment 1 that belief confidence constitutes a meaningful measure of value uncertainty, and having linked belief confidence with decision confidence, we then addressed another key question of this study: would people use their belief confidence to arbitrate between exploration and exploitation? In Experiment 2, the proportions of choice (75%) and rating trials (25%) were reversed to allow for more exploration, whereas in Experiment 1, people often gained a sufficient amount of information from the lower-value bandit simply through observing outcomes during the rating trials, thereby lowering the need for active exploration of that decision option during choice trials. Indeed, this new design largely increased the number of trials in which the participants chose the lower-value option: participants on average chose the subjectively perceived lower-value option on 22.1% of all trials as opposed to only 13.3% of all trials for the previous experiment—supporting the view that they were exploring more.

#### 2.2.1 Participants have meaningful insight into their gambling behavior

We first assessed whether decision confidence allowed any meaningful insight into people’s choice patterns, having now collected a considerably larger number of such choice trials. People made decisions on average 822 ms after the onset of the trial and chose the higher-value bandit—given previously observed outcomes—in on average 81.6% of all choice trials. Overall, people showed good resolution in their decision-confidence judgments, that is, their decision confidence varied with both the percentage of trials on which the higher-value option was chosen, as well as average reward points. Indeed, error rates (defined as the proportion of trials on which participants chose the lower-value option) differed reliably across confidence bins, F(1.7, 49.3) = 107.9, p <0.001, 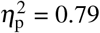, with a reliable linear trend, F(1, 29) = 140.9, p <0.001, 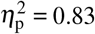, see Figure 4A. The same held for average rewards across decision-confidence quintiles, F(4, 116) = 117.0, p <0.001, 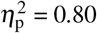, again with a reliable linear trend, F(1, 29) = 301.4, p <0.001, 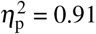, see Figure 4B.

**Figure 4.**
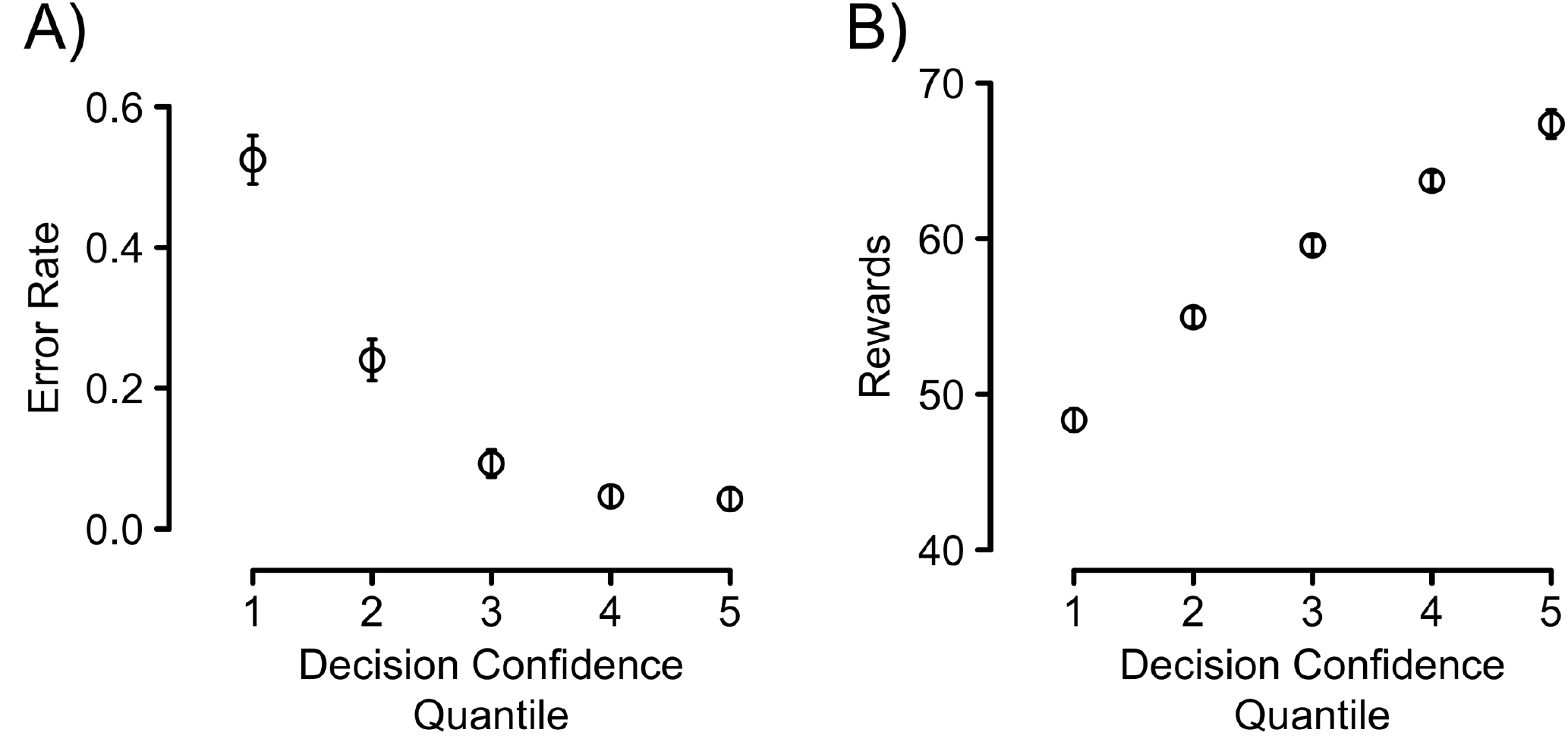
A) Error rates and B) average points won as a function of decision confidence. The data were binned according to decision confidence quintiles, which were formed within-subject. Errors are defined as trials on which people deviated from the ideal-observer model, that is trials on which they chose the arm of the bandit with so far the lower average in observed outcomes. All error bars are +/− 1 standard error of the mean for the respective y-axis values.

#### 2.2.2 Confidence-guided exploration

To test whether agents arbitrate between exploration and exploitation based on their belief confidence, we fitted a hierarchical, logistic regression model to participants’ choice-trial data. The model predicts the probability of choosing the lower-value option (exploration) from the belief confidence of the higher-value option, as well as the unsigned difference in value (DV) as a control variable, which we defined as abs(V_L_-V_R_). This model was identified as the best-fitting model from a formal model comparison approach based on *BIC* scores, which we report in the Supplementary Materials along with several other models. The Supplementary Materials furthermore include the results from a model that also includes belief confidence of the lower-value option as a (non-significant) predictor as well as a (significant) interaction between the two belief confidences (see also Figure 5A). However, this slightly more complex model did not fit the data as well (*BIC* = 11114.2) as the model presented here (*BIC* = 10923.9).

**Figure 5.**
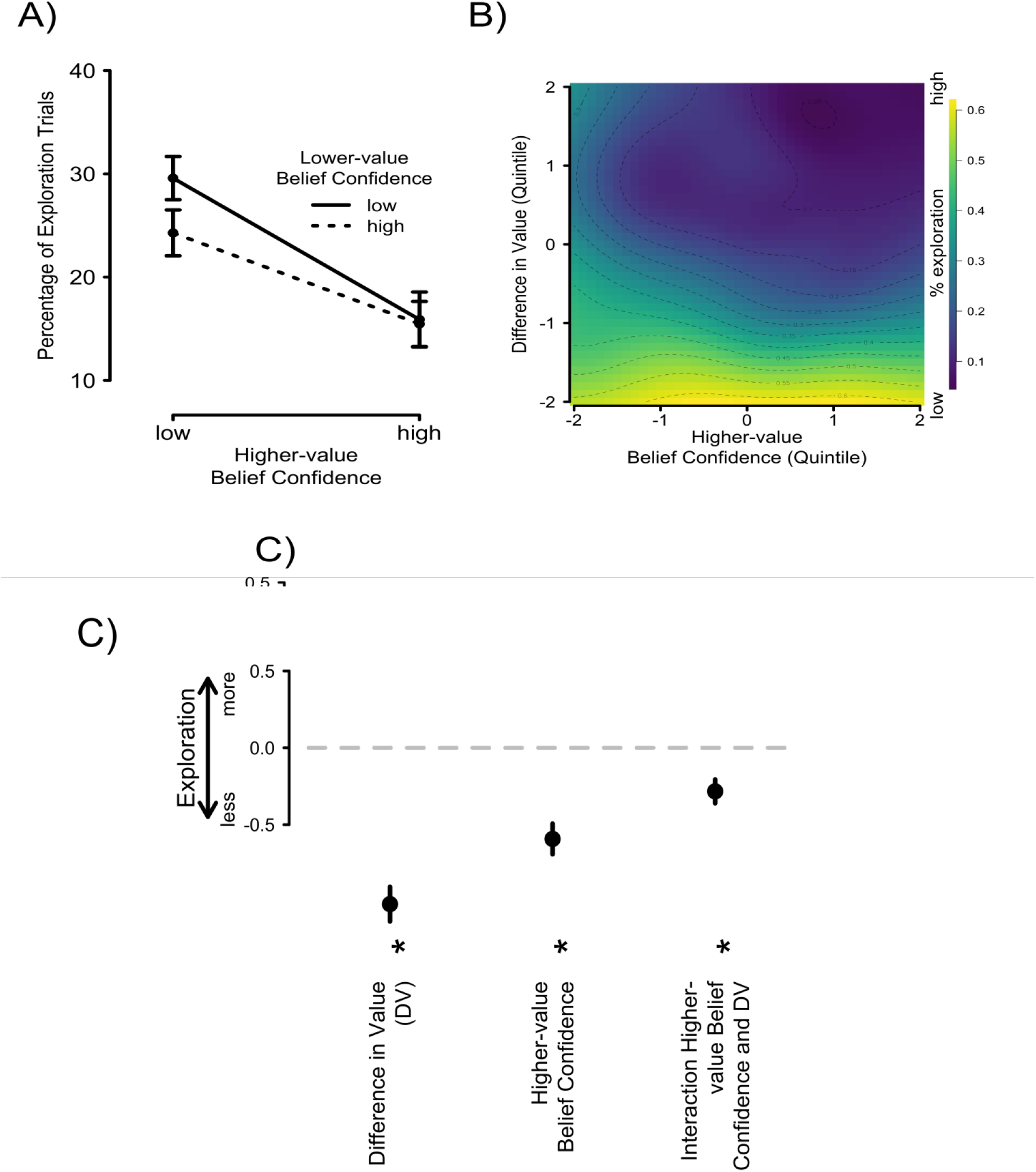
A) Proportion of trials in which participants chose the lower-value option (exploration), as a function of the belief confidence of the higher- and lower-value options. B) Proportion of trials in which participants chose the lower-value option (exploration trials), as a function of the belief confidence of the higher-value option and the difference in value. The dependent measure (exploration) is reflected in the color on the simulated grid, with lighter colors reflecting more exploration trials. C) Standardized, fixed regression coefficients from a mixed-model logistic regression model, predicting exploration. Positive, stronger parameter estimates reflect that an increase in this variable led to a larger tendency to explore. DV = difference in value. All error bars reflect +/− 1 standard error of the mean.

Critically, belief confidence of the higher-value option significantly predicted exploration, *β* = −0.59, p <0.001. This effect was negative, expressing that participants tended to explore more if their belief confidence was low, consistent with our prediction. The unsigned difference in value furthermore modulated choice significantly, *β* = −1.02, p <0.001, and negatively: the larger the absolute difference in value, the less participants chose to explore the lower-value option as arguably the overlap of the two value representations was small. Figure 5B presents how the difference in value (DV) and belief confidence of the higher-value choice option relate to exploration, with brighter colors reflecting higher proportions of exploratory choices. DV and confidence in the higher-value option did also interact reliably, *β* = −0.28, p <0.001. The standardized, fixed-effect regression coefficients are furthermore presented in Figure 5C. Together, these findings suggest that if the value representation of the preferred option is noisier, people tend to explore the inferior choice option more, compared to when it is precise. We thus conclude that belief uncertainty—as measured through belief confidence judgments—guides the trade-off between exploration and exploitation.

#### 2.2.3 Inter-individual differences in participants’ capability to track uncertainty

The increase in choice trials allowed us to obtain reliable measures of people’s metacognitive efficiency—the accuracy of their decision confidence judgments. The final question, which we aimed to address with our study was whether the degree by which people can accurately harness the level of uncertainty in their beliefs through confidence relates to their metacognitive efficiency, thus explaining some of the interindividual variability reported in the literature.

To estimate the former, we first fitted a hierarchical, linear regression model simultaneously to all participants’ rating-trial data to assess the degree to which belief-confidence estimates were driven by the variability of the past, observed outcomes or other control variables (mean of the past outcomes, the current trial within the block (log-transformed), and the arm of the bandit). Included in this model were two-way interactions between all variables except for the arm of the bandit, as well as the respective three-way interaction. A formal model comparison based on the *BIC* score (*BIC* = 17231.2), this model fitted the data best. A range of other models are presented in the Supplementary Materials.

The standardized regression weights for the fixed effects of all predictors are presented in Figure 6A. This simple, normative model of how beliefs should be formed over time constitutes an ideal observer model that can then be used to predict empirical responses. The influence of outcome variance on belief confidence was negative and reliably different from zero, *β*_sig_ = −0.37, p <0.001. The more variable past outcomes were, the less confident people became, consistent with our simple ideal-observer model. The mean of past outcomes had a weak positive, but also reliable influence on belief confidence, *β*_mu_ = 0.06, p = 0.04, the higher the outcomes people had observed for this arm, the more confident they judged their beliefs. The other two control variables, the current trial within the block (log-transformed), *β*_logtrial_ = −0.05, p = 0.46, and the arm of the bandit, *β*_trial_ = −0.02, p = 0.24, did not reliably predict belief confidence. Out of all two-way interactions, only the interaction between the standard deviation and mean of all previously observed outcomes was reliable, *β*_sig x mu_ = −0.14, p <0.001, as was the three-way interaction between standard deviation and mean of all previously observed outcomes and the current trial, *β*_sig x mu x logtrial_ = −0.04, p <0.01. None of the other interactions were reliable, abs(*β*s) <0.014, ps >0.54. Taken together, these findings suggest that people updated belief confidence similar to the a simple ideal-observer model. We then correlated the individual beta weights for the influence of outcome variance, *β*_sig_, with people’s metacognitive efficiency, log(M-ratio). We found that participants who were driven more in their belief-confidence estimates by the normative, ideal-observer belief confidence (outcome variance; i.e. larger negative regression weight for Beta Outcome *σ*) were more metacognitively efficient (higher metacognitive efficiency score), r = −0.37, p <0.05. In other words, people who were better at tracing the uncertainty present in the environment were better at distinguishing their own ‘good’ and ‘bad’ decisions, further linking the concepts of belief confidence and decision confidence. There was no such relationship between the beta weights for the influence of outcome mean, *β*_mu_, and metacognitive efficiency, r = 0.02, p = 0.94.

**Figure 6.**
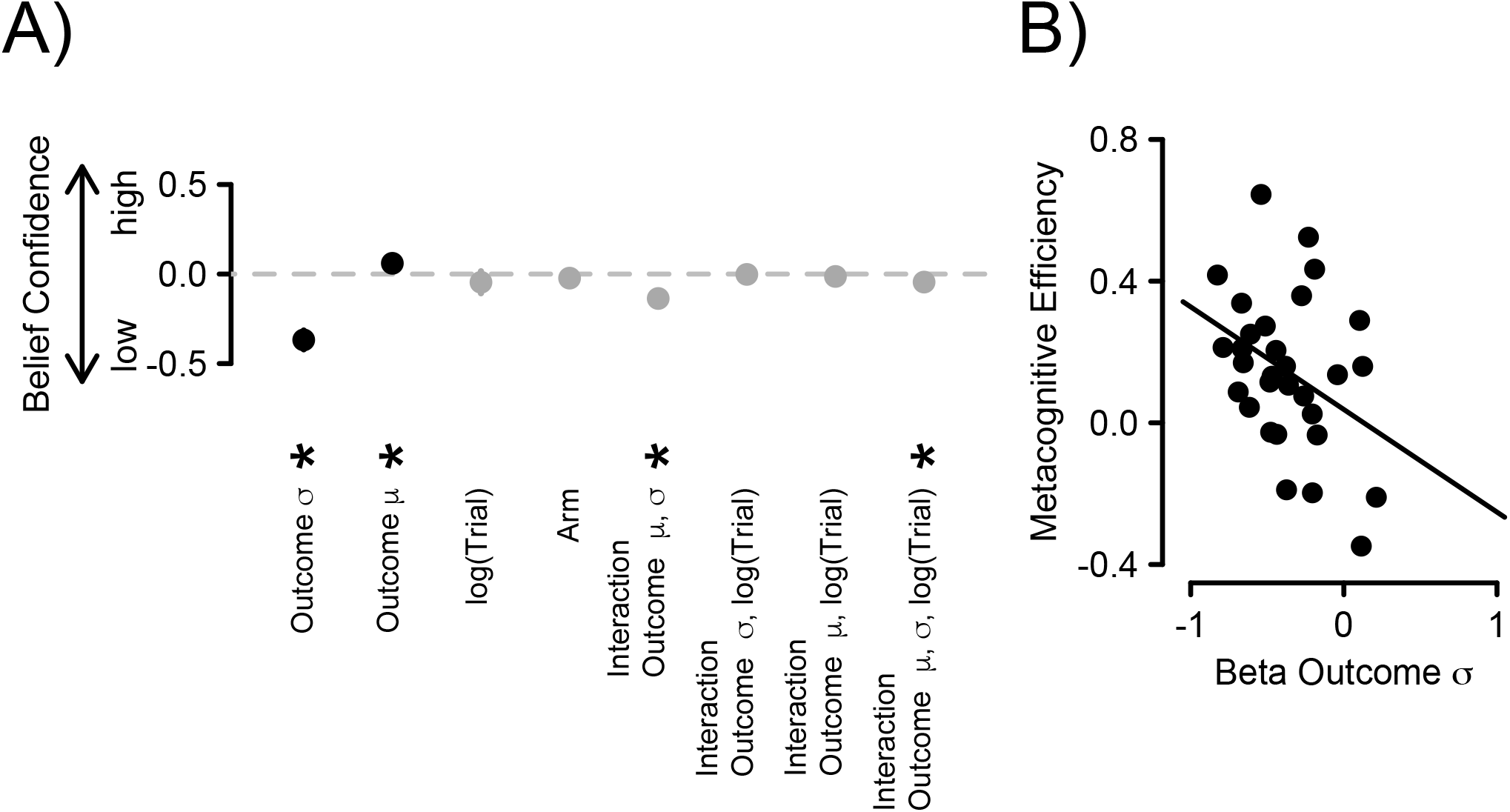
A) Standardized, fixed regression coefficients from a hierarchical, linear regression model. Positive parameter estimates reflect that an increment in this variable led to an increment in belief confidence. The error bars, which are almost entirely hidden behind the disks, reflect +/− 1 standard error of the mean. The black disks represent predictors linked to the observed outcomes, the grey disks represent control variables and interaction effects. B) Regression weights for the variance of past outcomes (ideal-observer model confidence) for each participant plotted against their metacognitive efficiency. *σ* = standard deviation; *μ* = mean.

The results of Experiment 2 suggest that people use their belief confidence to arbitrate between two extreme modes of behavior, exploration and exploitation. We moreover found that people whose belief confidence matched that of an ideal observer closer also showed better insight into their own decisions, thus suggesting a link between uncertainty in the belief representations (belief confidence) and decision confidence, further shedding light on the internal signals and cues that give rise to confidence in a value-based decision-making context.

## 3. Discussion

The present study used a novel variant of a bandit paradigm that allowed tracking of belief formation over time. Our key finding was that people rely on those internal estimates of uncertainty (belief confidence) to guide their behavior. This means that the confidence inherent in the representation of value beliefs can be used to arbitrate between exploration and exploitation. We found that noisy value representations—especially regarding the higher-value option—led participants to explore more. Such uncertainty-guided exploration matches and further extends previous findings of uncertainty-driven exploitation (e.g. Frank et al., 2009; Badre et al., 2012). The key difference, however, was that we have demonstrated that it is possible to directly measure the uncertainty that serves as a cue to action selection using confidence judgments. Previous studies, on the other hand, have focused entirely on computational estimates of uncertainty. Our findings are further strengthened by our novel, explicit measure of exploration. We define exploration as trials in which people chose the option they had previously rated as inferior. Formally, exploitation is defined as a state in which an agent is focussing its resources on the task or object that is currently believed to yield the highest payoff, whereas exploration is the expansion of the focus to search for other reward options (Sutton & Barto, 1998). In previous studies, exploitation has commonly been operationalized as the act of choosing the same option compared to the previous trial—whether this is referring to discrete choice options (e.g., Kolling et al., 2012) or different decision strategies that are combined continuously (e.g., Frank et al., 2009). However, such an operationalization is unable to capture whether the agent chooses the option that he perceives to currently yield the best outcomes (exploitation) or not (exploration). Here we defined as ‘exploration’ a situation in which people choose an option they explicitly rated as inferior.

Confidence-guided exploration takes its place alongside a range of other findings that suggest that confidence forms a cornerstone of cognitive control (for reviews see Yeung & Summerfield, 2012, 2014; Shea et al., 2014; Femandez-Duque et al., 2000; Nelson & Narens, 1990). For instance, research on metacognition in memory suggests that people use their internal, metacognitive signals to optimize learning, focusing on items they feel least confident about (Nelson & Leonesio, 1988), even when these metacognitive judgments were objectively wrong (Metcalfe & Finn, 2008; Hanczakowski, Zawadzka, & Cockcroft-McKay, 2014). Similarly, findings from the error-monitoring literature suggest that people commonly slow down after committing an error (Rabbitt, 1966), suggesting that they adopt a more accuracy-focused response regime to avoid further mistakes (Dutilh, Vandekerckhove, Forstmann, Keuleers, Brysbaert, & Wagenmakers, 2012). Relatedly, confidence has been shown to serve as an internal teaching signal to support learning (Guggenmos et al., 2016) and metacognition has been proposed as a mechanism to guide people’s decisions to cognitively offload by setting reminders to avoid memory failures (Risko & Gilbert, 2016). The notion that representations of uncertainty can be used by the brain to optimize behavior therefore extends and augments previous findings and discussions on metacognition.

We found that belief confidence predicted exploration in a linear way: the more uncertain people judged their value beliefs, the more likely they were to explore respective choice options. This finding is seemingly at odds with studies and theories that suggest confidence is related to curiosity in an inverse u-form shape (Kang, Hsu, Krajbich, Loewenstein, McClure, Wang, & Camerer, 2009; for a review of these findings see Butlin, 2010). These studies claim that when people feel least or most confident they are less likely to seek new information, whereas they are most curious for levels of medium confidence. However, it should be noted that these studies used a full confidence scale, that is a scale that reaches from 0% confidence (certainly wrong) over 50% confidence (guessing) to 100% confidence (certainly correct). In our study, belief confidence ratings were given on a scale ranging from 50-100% confidence. We chose this scale because we collected value belief and belief confidence ratings concurrently and therefore did not expect any error detection. These findings can thus be reconciled considering only the curiosity ratings for the upper half of the scale used in Kang et al. (2010).

Our study furthermore linked our new concept, belief confidence, to the more traditionally studied concept of decision confidence. We found that participants with more accurate insight into their decisions tended to be driven more by the variability in their experienced outcomes in their belief confidence. Relatedly, we found that belief confidence had an effect on how some people judged their choices. Critically, this was found not only for the chosen option (being certain regarding the higher value of the option participants selected increased their decision confidence) but for the belief confidence of the unchosen option: being certain about the value of the decision alternative people chose to forgo also increased their decision confidence. This finding fits and extends previous studies that reported that humans are capable of tracking counterfactual choice options (Boorman, Behrens, & Rushworth, 2011; see also Domenech & Koechlin, 2015).

The finding that the belief confidence of both choice options affects decision confidence furthermore extends and links to previous results from our lab. In an initial study, we found that confidence in a value-based binary decisions (i.e. decision confidence in the current framework) was associated with activity in both the vmPFC and the rlPFC (De Martino et al., 2013) in which the former tracked both difference in value estimates and confidence and the latter only confidence. In a subsequent study (De Martino et al., 2017), instead of using binary choice we elicited confidence into value estimates (i.e. belief confidence according to the taxonomy used here) we found again an involvement of vmPFC (expanding into dmPFC according to a functional gradient) but no confidence signal in rlPFC. In light of the results presented here, we are tempted to suggest a possible dissociation of roles these two regions might play in the readout of uncertainty. In the choice study (De Martino et al., 2013), participants repeatedly chose between different snack items presented on screen, reporting their decision confidence with every choice, whereas value beliefs and belief confidence were not measured. Given the tight link between belief confidence in the chosen item and decision confidence that we showed in the present study, it is possible that the vmPFC activations that were observed in the binary choice study (De Martino et al., 2013) are mainly reflections of belief confidence (i.e. the uncertainty into the value estimation). This possibility is consistent with a subsequent finding in which belief confidence was directly measured in an MRI study and in which activity in vmPFC was identified (De Martino et al., 2017; but see also Lebreton, Abitbol, Daunizeau, & Pessiglione, 2015). An intriguing possibility is that rlPFC is involved only in decision confidence (or the subsequent use of such for the purpose of cognitive control; cf. Badre et al., 2012), that is when participants are requested to explicitly report the probability of an action to be correct. This matches the notion that the rlPFC is involved in the readout of metacognitive reports (usually measured in choice and not in estimation tasks), and it is supported by findings that show that coupling strength of the rlPFC with vmPFC is predictive of how efficiently uncertainty is read out for the metacognitive report (De Martino et al., 2013), as well as that grey matter volume in this area correlates with participants’ metacognitive accuracy (Fleming et al., 2010; for a review of the role of rlPFC in metacognition also see Fleming & Dolan, 2012). This would suggest that low-level representations of uncertainty (measured by belief confidence here) are inherent to the computation performed by the cortical regions such as vmPFC for value beliefs (De Martino et al., 2013; 2017) or visual cortex for perceptual estimation (Fleming, Huijgen, & Dolan, 2012). However, following a choice, the uncertainty inherent in these low-level representations, together with uncertainty arising during the decision process such as response speed (Kiani et al., 2014) or familiarity and fluency (Koriat, 1997) can in turn inform metacognitive reports that are instantiated in rlPFC.

In the present study, we proposed that belief confidence arises from the belief updating process, both of which we measured using explicit ratings. Belief confidence is likely to reflect both the stochasticity inherent in the lotteries (risk), as well as epistemic uncertainty, which decreases gradually through learning. Relatedly, a recent study by Meyniel, Schlunegger, and Dehaene (2015a) proposed a learning paradigm that allows studying the development of confidence over time: Participants were presented with a sequence of one of two possible stimuli. From time to time, participants had to estimate the probability that they were currently in a hidden state, which generated these stimuli with different probabilities and their confidence regarding the correctness of their probability judgment. Meyniel and colleagues (2015a) found that people were able to trace the evolving transition probabilities and that their confidence judgments reflected the uncertainty inherent in such a learning process, stemming from both the inherent stochasticity of the task, insufficient exposure, as well as the sudden transitions of states that happened throughout the experiment, bearing some similarities to the concepts of expected and unexpected uncertainty (e.g. Yu & Dayan, 2005). The current study makes an important distinction between this concept of confidence (i.e. uncertainty in the judgement or belief confidence in our terminology) and the one that is usually reported in following a choice (i.e. the probability that an action is correct or decision confidence).

To conclude, our results provide a novel account of how uncertainty in value-belief judgments is constructed over time, and how people arbitrate between exploration and exploitation, with an uncertainty bonus biasing people towards exploration for the purpose of further information seeking (e.g. Daw et al., 2006; Frank et al., 2009). We suggest that our results carry implications for how the precision in the value representation for the chosen and unchosen option feeds into decision confidence, suggesting that uncertainty in the value-belief representations (belief uncertainty) also affects how confident people judge their choices, with higher decision confidence if decisions were based on precisely represented values of both the chosen and unchosen choice option. We showed that people who were better capable of tracing the uncertainty inherent in the environment also possessed higher metacognitive insight into their decisions. Our result therefore argue for a complementary role for these two types of confidence judgments.

## Acknowledgments

This study was funded by a Sir Henry Dale Fellowship (102612 /A/13/Z) awarded to Benedetto De Martino by the Wellcome Trust. We would like to thank Steve Fleming for his comments on a previous version of the manuscript and Jessica Hughes and Pradyumna Sepulveda for the help in the proofreading and editing of the manuscript.

## Supplementary Information

### Model Comparisons

#### Predicting Decision Confidence

To examine the effects of value beliefs and belief confidence on decision confidence, we compared several hierarchical regression models. A full description of these models can be found in Table S1, together with the resulting *BIC* scores in Figure S1. For the sake of simplicity, the main text presents only the best-fitting model (Model 8). Variables were included one by one and only kept in the model for the next step if they added value, that is if they reduced the *BIC*.

**Table S1.** Name and simplified formulas of the hierarchical regression models. Asterisks indicate interaction effects between variables. *ε* = error term.

| Model | Formula |
| --- | --- |
| Model 1 | $\text{DecConf} \sim \beta_0 + \beta_1[\text{Chosen Value}] + \varepsilon$ |
| Model 2 | $\text{DecConf} \sim \beta_0 + \beta_1[\text{Chosen Value}] + \beta_2[\text{Unchosen Value}] + \varepsilon$ |
| Model 3 | $\text{DecConf} \sim \beta_0 + \beta_1[\text{Chosen Value}] * \beta_2[\text{Unchosen Value}] + \varepsilon$ |
| Model 4 | $\text{DecConf} \sim \beta_0 + \beta_1[\text{Chosen Value}] * \beta_2[\text{Unchosen Value}] + \beta_3[\text{Chosen Belief Confidence}] + \varepsilon$ |
| Model 5 | $\text{DecConf} \sim \beta_0 + \beta_1[\text{Chosen Value}] * \beta_2[\text{Unchosen Value}] + \beta_3[\text{Chosen Belief Confidence}] + \beta_4[\text{Unchosen Belief Confidence}] + \varepsilon$ |
| Model 6 | $\text{DecConf} \sim \beta_0 + \beta_1[\text{Chosen Value}] * \beta_2[\text{Unchosen Value}] + \beta_3[\text{Chosen Belief Confidence}] * \beta_4[\text{Unchosen Belief Confidence}] + \varepsilon$ |
| Model 7 | $\text{DecConf} \sim \beta_0 + \beta_1[\text{Chosen Value}] * \beta_2[\text{Unchosen Value}] + \beta_3[\text{Chosen Belief Confidence}] + \beta_4[\text{Unchosen Belief Confidence}] + \beta_5[\text{Objective Accuracy}] + \varepsilon$ |
| Model 8 | $\text{DecConf} \sim \beta_0 + \beta_1[\text{Chosen Value}] * \beta_2[\text{Unchosen Value}] + \beta_3[\text{Chosen Belief Confidence}] + \beta_4[\text{Unchosen Belief Confidence}] + \beta_5[\text{Objective Accuracy}] + \beta_6[\log(\text{RT})] + \varepsilon$ |
| Model 9 | $\text{DecConf} \sim \beta_0 + \beta_1[\text{Chosen Value}] * \beta_2[\text{Unchosen Value}] + \beta_3[\text{Chosen Belief Confidence}] + \beta_4[\text{Unchosen Belief Confidence}] + \beta_5[\text{Objective Accuracy}] + \beta_6[\log(\text{RT})] + \beta_7[\log(\text{Trial Number})] + \varepsilon$ |
| Model 10 | $\text{DecConf} \sim \beta_0 + \beta_1[\text{Chosen Value}] * \beta_2[\text{Unchosen Value}] + \beta_3[\text{Chosen Belief Confidence}] * \beta_4[\text{Unchosen Belief Confidence}] + \beta_5[\text{Objective Accuracy}] + \beta_6[\log(\text{RT})] + \beta_7[\log(\text{Trial Number})] + \varepsilon$ |

**Figure S1.**
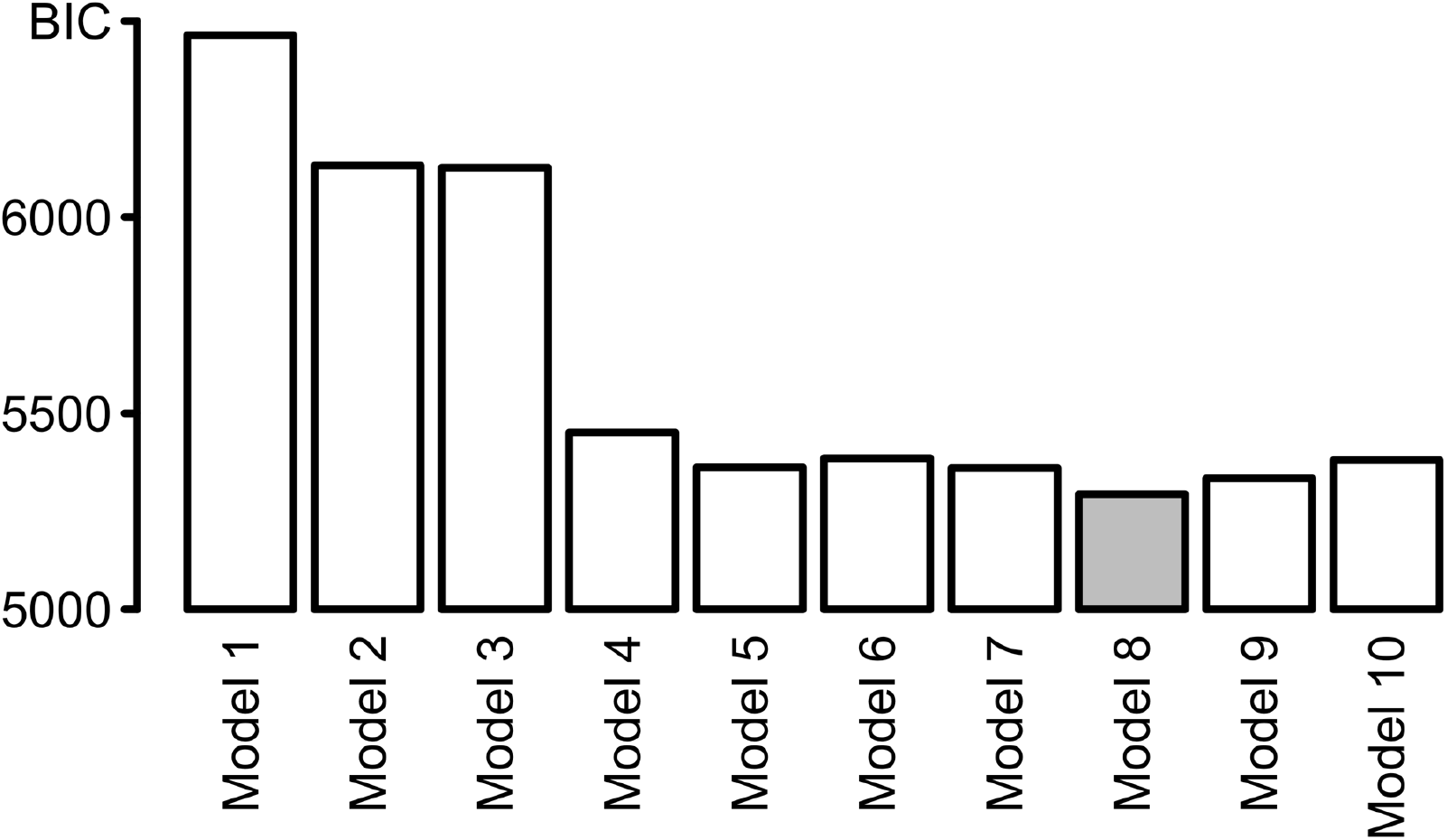
Resulting *BIC* values from model comparison approach from most parsimonious (Model 1) to most complex (Model 10). Model 8 (depicted in grey) was the best-fitting model and was reported in the main text for the sake of simplicity.

#### Predicting Exploration

Several hierarchical regression models were conducted to examine the effects of belief confidence on exploration. The full description of these models can be found in Table S2, the resulting *BIC* scores are presented in Figure S2A. For this analysis, variables were again included one by one and only kept in the model for the next step if they added value, that is if they reduced the *BIC*. The best-fitting model identified was Model 3, also presented in the main text. Figure S2B furthermore shows the regression weights for the most complex model, Model 5. This model differed from the best-fitting Model 3 in that it also included the lower-value belief confidence as a predictor for exploration, as well as the interaction of this additional variable with the other two regressors in the model. Replicating the results reported in the main text, belief confidence of the higher-value option significantly predicted exploration, *β* = −0.54, p <0.001. Again, this effect was found to be negative, reflecting that participants tended to explore more if their belief confidence was low. This relationship did not hold for the lower-value option, though, *β* = 0.08, p = 0.52. However, we found a reliable interaction between these factors, *β* = 0.21, p <0.01, reflecting that the belief confidence associated with the lower-value option affected exploration only if the belief confidence associated with the higher-value option was low (see also Figure 5A in the main text). The unsigned difference in value furthermore modulated choice significantly, *β* = −1.02, p <0.001, and negatively: The larger the absolute difference in value, the less participants chose to explore the lower-value option as arguably the overlap of the two value representations was small. Again, this finding replicated the results reported in the main text. DV and confidence in the higher-value option did also again interact reliably, *β* = −0.20, p = 0.04. None of the other interaction terms were reliable, abs(*β*s) <0.09, ps >0.20.

**Table S2.** Name and simplified formulas of the hierarchical regression models. Asterisks indicate interaction effects between variables. *ε* = error term.

| Model | Formula |
| --- | --- |
| Model 1 | $\text{DecConf} \sim \beta_0 + \beta_1[\text{Higher-value Belief Confidence}] + \varepsilon$ |
| Model 2 | $\text{DecConf} \sim \beta_0 + \beta_1[\text{Higher-value Belief Confidence}] + \beta_2[\text{Difference in Value}] + \varepsilon$ |
| Model 3 | $\text{DecConf} \sim \beta_0 + \beta_1[\text{Higher-value Belief Confidence}] * \beta_2[\text{Difference in Value}] + \varepsilon$ |
| Model 4 | $\text{DecConf} \sim \beta_0 + \beta_1[\text{Higher-value Belief Confidence}] * \beta_2[\text{Difference in Value}] + \beta_3[\text{Lower-Value Belief Confidence}] + \varepsilon$ |
| Model 5 | $\text{DecConf} \sim \beta_0 + \beta_1[\text{Higher-value Belief Confidence}] * \beta_2[\text{Difference in Value}] * \beta_3[\text{Lower-Value Belief Confidence}] + \varepsilon$ |

**Figure S2.**
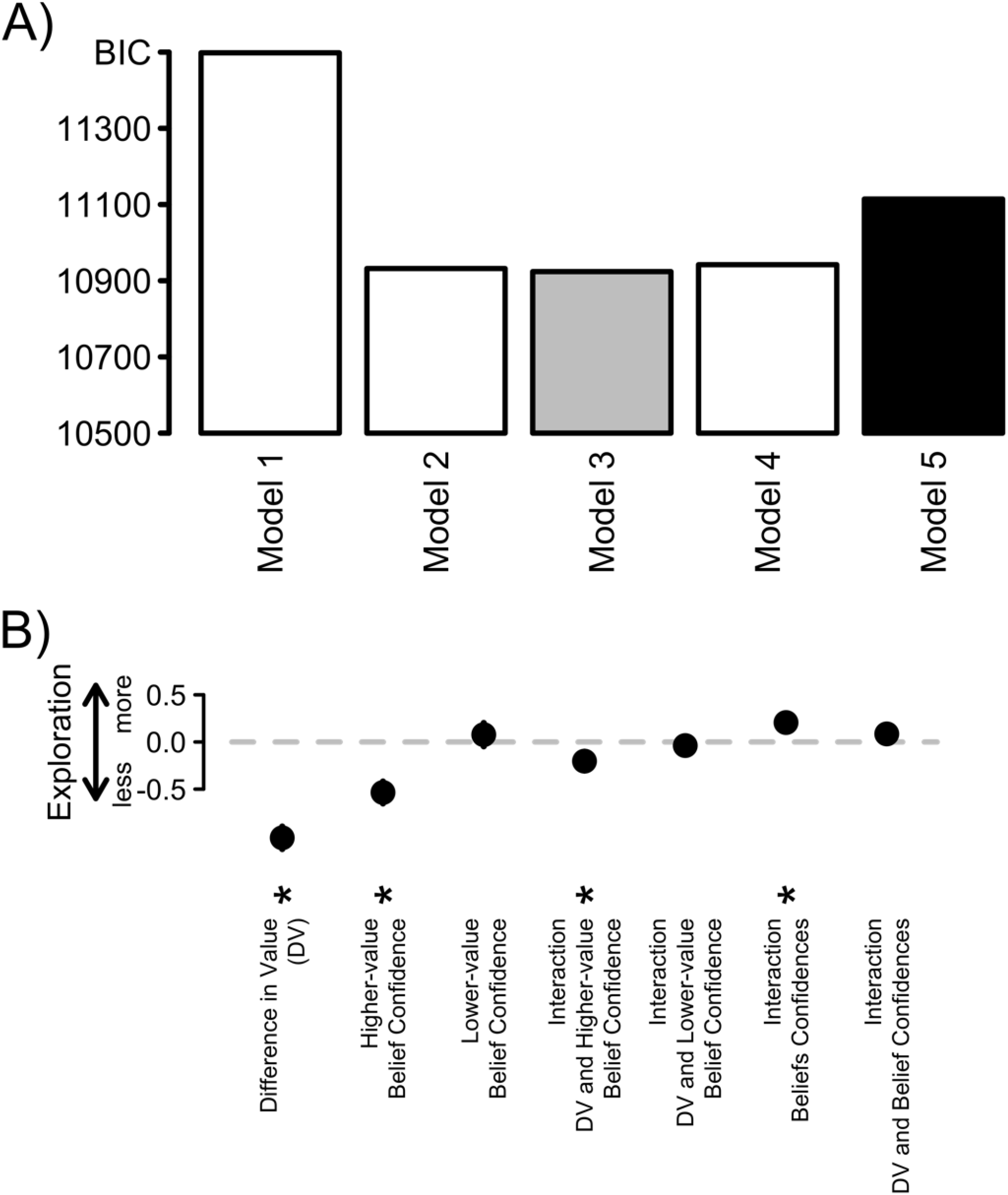
A) Resulting *BIC* values from model comparison approach from most parsimonious (Model 1) to most complex (Model 5). Model 3 (depicted in grey) was the best-fitting model and reported in the main text. The standardized, fixed regression coefficients resulting from Model 5 (depicted in black) are presented in B). Positive, stronger parameter estimates reflect that an increase in this variable led to a larger tendency to explore. DV = difference in value. All error bars reflect +/− 1 standard error of the mean.

#### Predicting Belief Confidence

We compared several hierarchical regression models to assess the influence of the objective variance and mean of previously observed outcomes. Table S3 contains a full description of these models, together with their resulting *BIC* scores in Figure S3. Variables were included one by one and only kept in the model for the next step if they added value, that is if they reduced the *BIC*. The model from the main text, the best-fitting model in terms of *BIC*, is referred to as Model 6 here.

**Table S3.** Name and simplified formulas of the hierarchical regression models. Asterisks indicate interaction effects between variables. *ε* = error term.

| Model | Formula |
| --- | --- |
| Model 1 | $\text{BelConf} \sim \beta_0 + \beta_1[\text{Outcome Variance}] + \varepsilon$ |
| Model 2 | $\text{BelConf} \sim \beta_0 + \beta_1[\text{Outcome Variance}] + \beta_2[\text{Outcome Mean}] + \varepsilon$ |
| Model 3 | $\text{BelConf} \sim \beta_0 + \beta_1[\text{Outcome Variance}] * \beta_2[\text{Outcome Mean}] + \varepsilon$ |
| Model 4 | $\text{BelConf} \sim \beta_0 + \beta_1[\text{Outcome Variance}] * \beta_2[\text{Outcome Mean}] + \beta_3[\log(\text{Trial Number})] + \varepsilon$ |
| Model 5 | $\text{BelConf} \sim \beta_0 + \beta_1[\text{Outcome Variance}] * \beta_2[\text{Outcome Mean}] * \beta_3[\log(\text{Trial Number})] + \beta_4[\text{Arm}] + \varepsilon$ |
| Model 6 | $\text{BelConf} \sim \beta_0 + \beta_1[\text{Outcome Variance}] * \beta_2[\text{Outcome Mean}] * \beta_3[\log(\text{Trial Number})] + \beta_4[\text{Arm}] + \varepsilon$ |
| Model 7 | $\text{BelConf} \sim \beta_0 + \beta_1[\text{Outcome Variance}] * \beta_2[\text{Outcome Mean}] * \beta_3[\log(\text{Trial Number})] * \beta_4[\text{Arm}] + \varepsilon$ |

**Figure S3.**
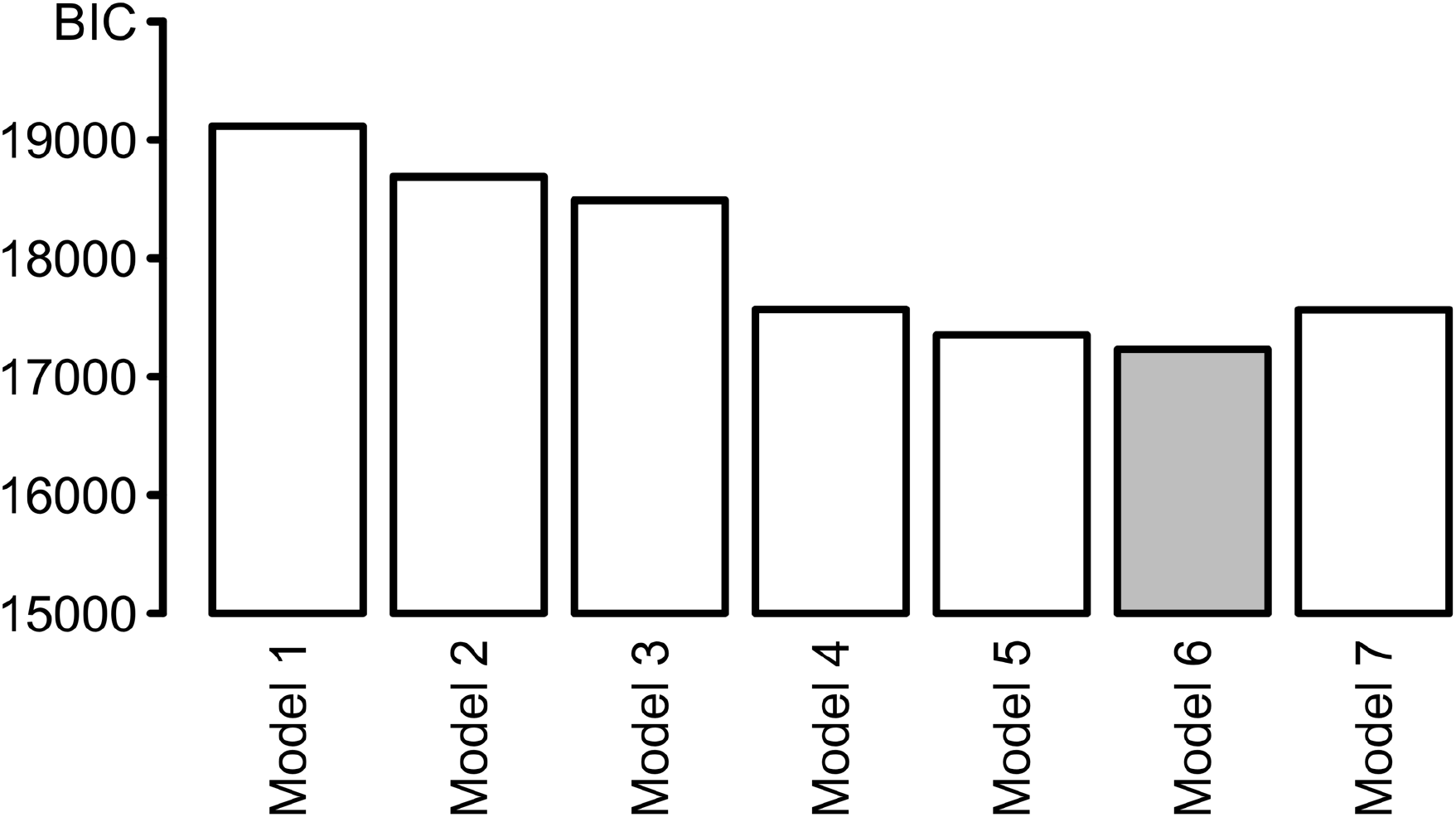
Resulting *BIC* values from model comparison approach from most parsimonious (Model 1) to most complex (Model 7). Model 6 (depicted in grey) was the best-fitting model and presented in the main text.

